# A mathematical model for phenotypic heterogeneity in breast cancer with implications for therapeutic strategies

**DOI:** 10.1101/2021.06.04.447174

**Authors:** Xin Li, D. Thirumalai

## Abstract

Inevitably, almost all cancer patients develop resistance to targeted therapy. Intratumor heterogeneity is a major cause of drug resistance. Mathematical models that explain experiments quantitatively are useful in understanding the origin of intratumor heterogeneity, which then could be used to explore scenarios for efficacious therapy. Here, we develop a mathematical model to investigate intratumor heterogeneity in breast cancer by exploiting the observation that HER2+ and HER2− cells could divide symmetrically or asymmetrically. Our predictions for the evolution of cell fractions are in quantitative agreement with single-cell experiments. Remarkably, the colony size of HER2+ cells emerging from a single HER2− cell (or vice versa), which occurs in about four cell doublings, also agrees with experimental results, without tweaking any parameter in the model. The theory quantitatively explains experimental data on the responses of breast tumors under different treatment protocols. We then used the model to predict that, not only the order of two drugs, but also the treatment period for each drug and the tumor cell plasticity could be manipulated to improve the treatment efficacy. Mathematical models, when integrated with data on patients, make possible exploration of a broad range of parameters readily, which might provide insights in devising effective therapies.

## INTRODUCTION

Nearly 10 million people died of cancer worldwide in 2020^1^, despite innovations in the development of many novel drugs. In principle, the advent of new technologies ought to make drugs highly efficacious while minimizing toxicity. The next-generation sequencing allows us to design personalized therapy, targeting specific genetic variants which drive disease progression^2,3^. However, drug resistance ultimately occurs, regardless of targeted therapeutic protocols, which poses a formidable challenge for oncologists^4^. A deeper understanding of the underlying resistance mechanism could be useful in controlling the tumor burden and its relapse.

Intratumor heterogeneity, which denotes the coexistence of cancer cell subpopulations with different genetic or phenotypic characteristics in a single tumor^5,6^, is the prominent cause of drug resistance and recurrence of cancers^7–9^. With the development of deepsequencing technologies and sequencing at the single cell level^10,11^, intratumor *genetic* heterogeneity has been observed in many cancer types^12–17^. Meanwhile, increasing evidence shows that *phenotypic* variations in tumor cells (without clear genetic alterations) also play a crucial role in cancer development, and is presumed to be one of the major reasons for the development of drug resistance in cancer therapy^7,18^. However, the underlying mechanism of intratumor heterogeneity induced by the phenotypic variability of cancer cells is still elusive, which represents an obstacle for the development of efficient treatments for cancer patients^19^.

The phenotypic heterogeneity of normal cells can emerge from cellular plasticity, which is the ability of a cell to adopt different identities. Cellular plasticity is widespread in multicellular organisms, dictating the development of organism, wound repair and tissue regeneration^20–22^. One of the best known examples is the differentiation hierarchies in stem cells, which leads to the production of progenitor cells, followed by the mature differentiated cells^23,24^.

It has been proposed that cancer might be derived from cancer stem (or initiating) cells. The cancer stem cells are similar to normal stem cell, but possess the ability to produce all cell types found in a tumor sample, resulting in intratumor heterogeneity^25–27^. However, the prospects of a hierarchical organization, and also the unidirectional differentiation of cancer stem cells have been challenged by recent experimental observations^28–31^. Some ‘differentiated’ cancer cells are capable of switching back to the cancer stem cells in breast cancer^28,29^. Melanoma cells do not show any hierarchically organized structure as cells are capable of switching between different phenotypes reversibly^30,31^. Several models that assume reversible state transitions have been proposed to explain the observed stable equilibrium among cancer cell subpopulations with different phenotypes^28,32^. However, a detailed understanding of the underlying mechanism driving the cell state transition is still lacking, as most previous experimental observations are based on measurements from bulk cell populations^28,29,31^.

A recent insightful experiment tracked the evolution of a single circulating tumor cell derived from estrogen-receptor (ER)-positive/human epidermal growth factor receptor 2 (HER2)-negative (ER+/HER2−) breast cancer patients *in vitro*^33^. Surprisingly, HER2+ cells (with expression of HER2) emerge from a cell colony grown from a single HER2− cell within four cell doublings and vice versa. The single-cell level experiment demonstrates that reversible transitions occurred between the two breast cancer cell types, thus providing a clue to understanding the nature of cancer cell plasticity observed in this and other experiments^28,29,31,33^. Because normal stem cell can differentiate into non-stem cells through asymmetric cell division^23^, it is possible that cancer cells might also change their identity by asymmetric division^34^, which is a potential cause of intratumor heterogeneity.

We noticed that the emergence of an altered cell phenotype is to be coupled to cell division, as indicated by the experiments that a cell of a specific genotype produces daughter cells with an altered phenotype^33^. We developed a theoretical model to describe the establishment of intratumor heterogeneity from a single type of breast circulating tumor cells. In quantitative agreement with experiments, our model captures the tumor growth dynamics under different initial conditions. It also naturally explains the emergence and evolution of intratumor heterogeneity, initiated from a single cell type, as discovered in a recent experiment^33^. Based on the parameter values derived from the tumor growth experiments, we predict the evolution of cell fractions, and the colony size for the appearance of HER2+ (HER2−) cell types starting from a single HER2− (HER2+) cell. Remarkably, the predictions agree well with the experimental observations. As a consequence of intratumor heterogeneity, drug resistance develops rapidly, which we also reproduce quantitatively. By exploring a range of parameters in the mathematical model, we found that several factors strongly influence the growth dynamics of the tumor. The insights from our study may be useful in devising effective therapies^33,35^.

## MODEL AND METHODS

### Brief summary of the circulating breast cancer cells experiments^33^

Using microfluidic CTC-iChip and Fluorescence-activated cell sorting (FACS), two different types of tumor cells, HER2 positive (HER2+) and HER2 negative (HER2−) cells, are extracted and separated from fresh whole blood of patients originally diagnosed with HER2− breast cancer. No genetic mutation has been identified between the two types of cells through single-cell sequencing. A heterogeneous cell population is observed in a few weeks as a cell colony (or a single cell) of either type only (tagged with fluorescent proteins initially) grows in an ultra-low attachment plate with tumor sphere medium (Roswell Park Memorial Institute (RPMI) 1640 Medium, epidermal growth factor (20 ng/ml), basic fibroblast growth factor (20 ng/ml), 1X B27, 1X antibiotic/antimycotic) under hypoxic (4% *O*_2_) conditions.

For mouse xenograft assays and drug treatment, 6-week-old female NOD scid gamma (NSG) mice from Jackson Laboratories were used. The green fluorescent protein-luciferase labelled circulating tumor cells were injected into the fourth right mammary fat pad. The growth of tumors in the mice was tracked weekly using IVIS Lumina II (PerkinElmer) for in vivo imaging.

### Model

A recent experiment shows^33^ that HER2+ (HER2−) cells can produce daughter cells of the other type, HER2− (HER2+) cells. In addition, a direct observation of the coupling between cell phenotype changes and cell divisions is reported in other related experiments^34^. We developed a mathematical model in which the cell plasticity is coupled to cell division, as illustrated in Fig. **1a** to understand the reversible transitions between different cell types, the establishment of intratumor heterogeneity, and also the complex drug responses observed in experiments. The HER2+ (HER2−) cells divide in a symmetric way producing two identical HER2+ (HER2−) cells with a rate *K*_1_ (*K*_2_) (see the green arrows in Fig. 1**a**). Asymmetric cell division can also occur for HER2+ (HER2−) cells generating one HER2+ and one HER2− cell with a rate *K*_12_ (*K*_21_) (see the blue arrows in Fig. 1**a**), as noted in a similar experiment for breast cancer cells^34^. It is also possible to produce two identical cells which differ from the parent cell type through another type of symmetric cell division with a rate *K*_13_ (*K*_31_)^36^ (see the grey dotted arrows in Fig. 1**a**). Because such an event is rare in the experiments^33^ we do not consider it. Therefore, instead of a unidirectional process for the stem cell differentiation^37^, we introduced a bidirectional state transition model through cell divisions without the hierarchical organization^28,31^.

**Figure 1:**
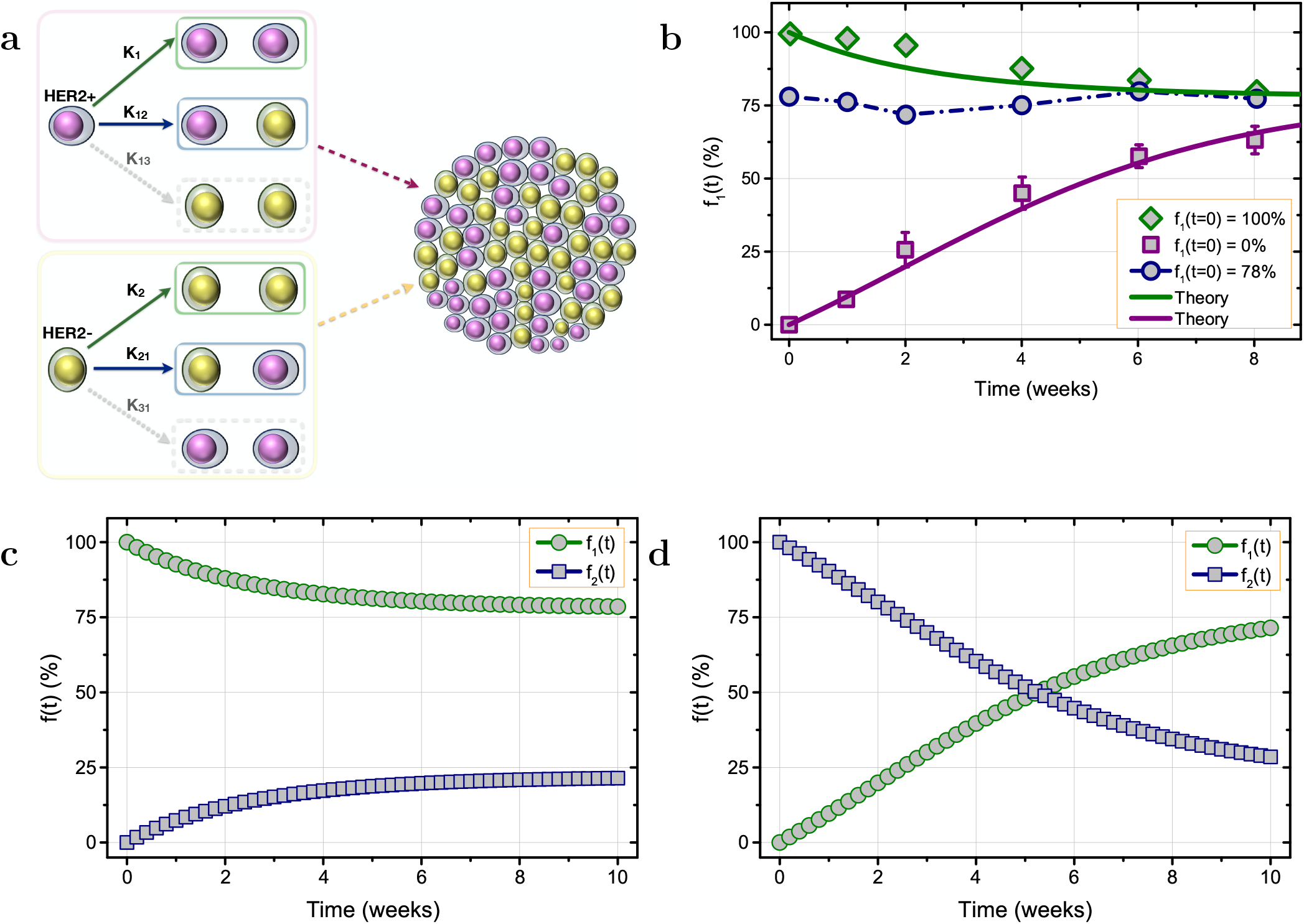
The dynamics of HER2+/HER2− cells. (**a**) Illustration of the intratumor heterogeneity model for breast cancer. Both HER2+ and HER2− breast circulating tumor cells may divide symmetrically, producing two identical HER2+ and HER2− cells with rates *K*_1_ and *K*_2_, respectively. They can also divide in an asymmetric manner by producing one HER2+ and one HER2− cell with rates *K*_12_ and *K*_21_. The two cell types could divide symmetrically but produce the other cell type (see the processes with rates of K13 and K31). A heterogeneous cell colony composed of both HER2+ and HER2− cells is established, irrespective of the initial cell states. (**b**) Experimental data for the fraction (*f*_1_(*t*)) of HER2+ cells as a function of time for three initial conditions: starting with HER2+ cells only (symbols in green), HER2− cells only (symbols in violet), and the parental cultured circulating tumor cells (symbols in navy). Theoretical predictions are shown by the solid lines. The dash dotted line for the case of parental cultured circulating tumor cells is to guide the eye. The time course of HER2+ (green) and HER2− (navy) cell fractions (*f*_1_(*t*), and (*f*_2_(*t*))) under the initial condition, (**c**) *f*_1_(*t* = 0) = 1, and (**d**) *f*_1_(*t* = 0) = 0.

The evolution of the population size *N*_1_(*t*) for HER2+ cells and *N*_2_(*t*) for HER2− cells are described by the following coupled equations,

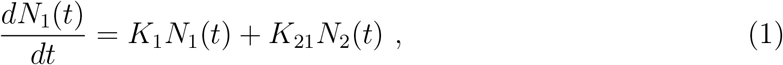

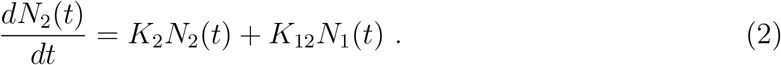

The total population is *N* (*t*) = *N*_1_(*t*) + *N*_2_(*t*). Several assumptions are used in our model. First, we do not consider apoptosis explicitly (unless drug treatments are applied), which, if needed, can be incorporated into the rate constants *K*_1_ and *K*_2_, allowing us to consider them as effective growth rates. We also neglected the symmetric division events (*K*_13_*, K*_31_ in Fig. 1**a**) that would produce two identical cells of a different type from their mother cell because it rarely occurs in the experiments and can be simply integrated into the second terms in Eqs. (1) and (2) in further studies. In addition, the carrying capacity is not reached in the experiments (see Fig. 3 and also Fig. S1 in the Supplementary Information that a rapid tumor growth is still observed at the end of experiments), so we did not use logistic differential equations.

Let us define the fraction of HER2+ cell in the whole population as *f*_1_(*t*) ≡ *N*_1_(*t*)*/N* (*t*), and the fraction *f*_2_(*t*) of HER2− is given by *f*_2_(*t*) = 1 − *f*_1_(*t*). Then, the evolution of *f*_1_(*t*) can be derived from Eqs. (1) and (2), leading to

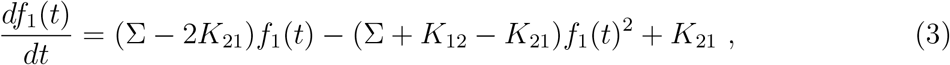

with Σ ≡ *K*_1_ − *K*_2_. Eq. (3) shows that the evolution of the cell fraction *f*_1_(*t*) (*f*_2_(*t*)) only depends on the rate difference Σ of the two symmetric cell divisions but not their absolute values *K*_1_ and *K*_2_. Therefore, there are only three (*K*_12_, *K*_21_ and Σ) free parameters in the model.

For simplicity, we assume that *K*_12_ = *K*_21_ ≡ *K*_0_ for the production of one cell type from the other. Actually, we find that these two rates are quite small as the other cell type appears only after several rounds of cell division events in the single cell experiments, and it is not necessary to give different values to them in order to explain all the experimental results discussed in this article. This turns out to be a reasonable assumption. Then, Eq. (3) can be further simplified as

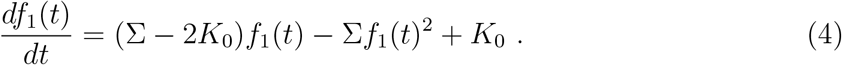

## RESULTS

### Phenotypic equilibrium in a heterogeneous cancer cell population

To demonstrate the cellular plasticity, a colony of a single cell type (either HER2+ or HER2− cells) is grown in culture for eight weeks in the experiments (see the Model and Methods for experimental details)^33^. Surprisingly, HER2− (HER2+) cells, naturally emerge from the initial HER2+ (HER2−) cell seeding within four weeks. The time course of the HER2+ cell fraction, *f*_1_(*t*), is shown in Fig. 1**b** for different initial conditions. The fraction *f*_1_(*t*) decreases slowly, reaching a plateau with *f*_1_ ≈ 78% after eight weeks of growth (see the green diamonds in Fig. 1**b**) starting exclusively from HER2+ cells. On the other hand, *f*_1_(*t*) increases to 63% (not reaching a plateau yet, a longer time required to reach the steady states) from zero rapidly during the same time period, if the cell colony is seeded only from HER2− cells (see the violet squares in Fig. 1**b**). Finally, the HER2+ cell faction, *f*_1_(*t*), almost does not change with time if the initial population is a mixture of both cell types derived from the parental cultured circulating tumor cells directly (see the navy circles in Fig. 1**b**). Therefore, a steady state level (with *f*_1_ ≈ 78%, the value in the parental cultured circulating tumor cells) is established between the two different cell phenotypes at long times, irrespective of the initial cell fraction.

To understand the experimental findings summarized in Fig. 1**b**, we developed a mathematical model as illustrated in Fig. 1**a** and described in the Model and Methods section. From Eq. (4) above and the stable equilibrium condition observed for the two cell populations in experiments we obtain,

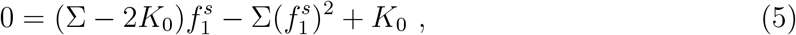

with the HER2+ cell fraction 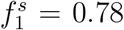 in the stationary state. Hence, we only need one more equation to fix the two free parameters (*K*_0_ and Σ) in Eq. (5).

Given the initial condition, *f*_1_(*t* = 0) = 0, we find that 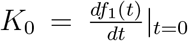 from Eq. (4) directly. Therefore, the parameter value *K*_0_ ≈ 0.09 per week is obtained using the first two data points from the experiments starting with only HER2− cells (see the violet squares in Fig. 1**b**). Finally, the value of Σ can be calculated from Eq. (5), which leads to Σ ≈ 0.3 given the stable equilibrium condition 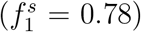 found in the two cell populations in experiments (see Fig. 1**b**). Hence, the time course of *f*_1_(*t*) can be calculated by solving Eq. (4), given any initial condition, *f*_1_(*t* = 0). Our theoretical predictions agree quantitatively with experiments (see the green, violet solid lines and the diamonds, squares in Fig. 1**b**), which is interesting considering that we only used two experimental data points. The time course of *f*_1_(*t*) and *f*_2_(*t*) (the fraction of HER2− cell) in the same plot under two initial conditions (see Fig. 1**c** and Fig. 1**d**) shows that the cell fraction conversion from HER2+ to HER2− is very slow. In contrast, the reverse process is rapid (see the slopes of the curves in Fig. **1c** and Fig. 1**d**). However, it takes shorter time for the system to reach the stationary state in the former case due to the large value of 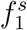 in experiments.

### Growth dynamics of cancer cell populations

The circulating tumor cells of HER2+ have a higher proliferation rate compared to HER2−, as noted both in *in vitro* and *in vivo* experiments (see Fig. S1 in the SI, and it is also supported by the experiments showing a much higher expression of the proliferation marker Ki67 in HER2+ cells compared to HER2− cells)^33^. It is consistent with the predictions of our model, which shows that the rate difference, Σ ≡ *K*_1_ − *K*_2_ ≈ 0.3, between the two cell types. Combined with the assumption that *K*_12_ = *K*_21_ ≡ *K*_0_, it also explains both the fast increase in *f*_1_(*t*) for the case when growth is initiated from HER2− cells, and the slowly decay of *f*_1_(*t*) as initial condition is altered (Fig. 1**b**). The different dynamics of HER2+ cell is also associated with it being a more aggressive phenotype, including increased invasiveness, angiogenesis and reduced survival^38,39^.

To understand the growth dynamics of the cell populations as a function of initial conditions (Fig. S1) quantitatively, we need to determine either *K*_1_ or *K*_2_. The other rate constant can be calculated using, *K*_1_ − *K*_2_ ≈ 0.3. We tuned the value of *K*_2_ and calculated the total tumor size from Eqs. (1)–(2) directly. Then, we compared our theoretical calculations to the experimental results to find the optimum value of *K*_2_ ≈ 0.7 per week which captures the tumor growth (see the green, navy solid lines and symbols in Fig. S1 in the SI). Note that *K*_2_ ≈ 0.7 implies that *K*_1_ ≈ 1.0 per week. We can also predict the growth dynamics at different initial conditions, which could be tested in similar experiments. From the values of the rate constants, we would expect that the frequency for symmetric cell division (the two daughter cells are identical to the parent cell) is much higher than the asymmetric case for both the cell types (*K*_1_ > *K*_2_ ≫ *K*_12_, *K*_21_). This prediction could be tested using single cell experiments.

### Cancer cell plasticity observed in single cell experiments

To further validate the model, we calculated the percentage of HER2+/HER2− cells as a function of the cell colony size starting from a single HER2+ or HER2− cell. The sizes of the cell colonies have been measured in experiments (see the histograms in Fig. 2)^33^. From Eqs. (1)–(2), we computed the HER2+ (HER2−) cell fraction, *f*_1_(*f*_2_), as a function of the cell colony size *N* with the initial conditions, *N*_1_(*t* = 0) = 1 and *N*_2_(*t* = 0) = 0 (*N*_1_(*t* = 0) = 0 and *N*_2_(*t* = 0) = 1) using the same parameter values as given above. Our theoretical predictions (see the solid line in Figs. 2**a** and 2**b**) capture the main features of the experimental observations without adjusting any parameter, especially for Fig. 2**a**. We also found that the HER2− cell fraction (*f*_1_) decreases faster than the HER2+ cell fraction (*f*_2_) as a function of the colony size (*N*), which is due to the higher symmetric division rate (*K*_1_ > *K*_2_) of HER2+ cells (Fig. 1**c** and 1**d**).

**Figure 2:**
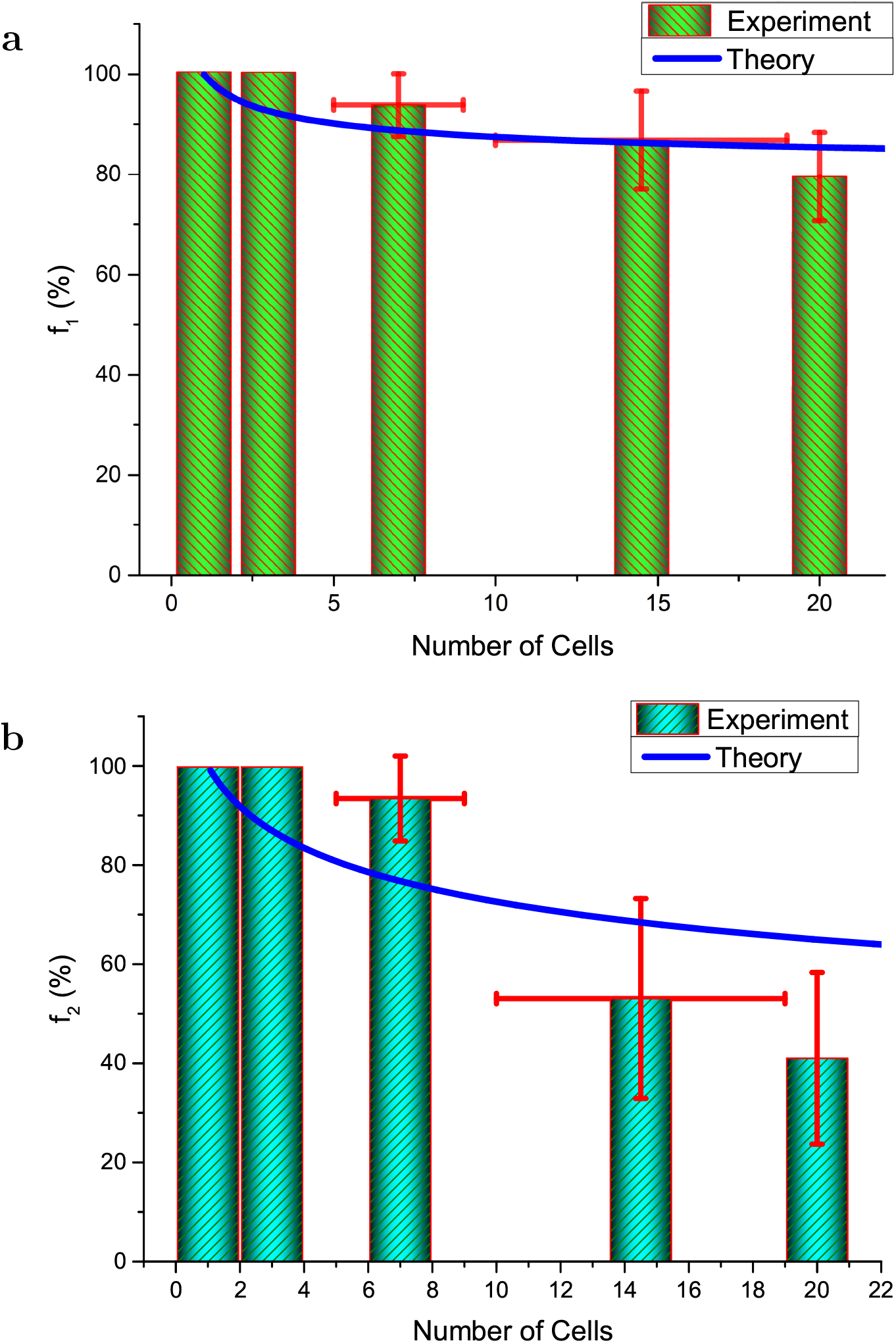
The interconversion between HER2+ and HER2− cell types. (**a**) The HER2+ cell fraction, *f*_1_ (percentage), as a function of the total population size *N* in a colony grown from a single HER2+ cell. (**b**) The HER2− cell fraction, *f*_2_ (percentage), as a function of *N* as the system develops from a single HER2− cell. The error bar in y-axis gives the standard variation, while the error bar in x-axis indicates the cell number range in which the cell fraction is calculated.

Similarly, from Eqs. (1)–(2), we can calculate the total number of cells N when the number of the other cell type *N*_1_ (*N*_2_) equal to one starting from *N*_2_ = 1(*N*_1_ = 1). The value of *N* is around 5 and 8 obtained from our model for HER2+ and HER2− cells, respectively. And the experimental values are found to be 5 to 9 cells, which agrees well with our theoretical predictions. Therefore, the model explains the experimental observation that one cell phenotype can emerge from the other spontaneously after four cell divisions.

### Drug response in a heterogeneous breast cancer cell population

It is known that HER2+ cells appear in patients initially diagnosed with ER+/HER2− breast cancer during treatment^40,41^. Although each cell subpopulation is sensitive to a specific drug, the heterogeneous tumor shows varying responses for distinct treatment protocols (see Fig. 3 as an example). The size of an untreated tumor increases rapidly (see the green circles) initiated from a mixture of two cell types. A clear response is noted when Paclitaxel (targeting HER2+ cells) is utilized, which results in reduced tumor growth (see the navy down triangles). Surprisingly, the tumor continues to grow rapidly, with no obvious response, if treated by Notch inhibitor (see the dark yellow squares). This is unexpected as the growth of HER2− cells (sensitive to Notch inhibitor) is supposed to be inhibited by the drug. Finally, the combination therapy with both the drugs, Paclitaxel and Notch inhibitor, administered to the tumors simultaneously effectively delays the tumor recurrence (see the violet up triangles). Given the potential synergies between these drugs to constrain the growth of heterogeneous populations of cancer cells, a deeper understanding of the drug resistance mechanism, and evolutionary dynamics of each subpopulation quantitatively is warranted. Here, we use the theoretical model above (see Fig. 1**a**) to explore diverse responses under different drug treatments (Fig. 3).

**Figure 3:**
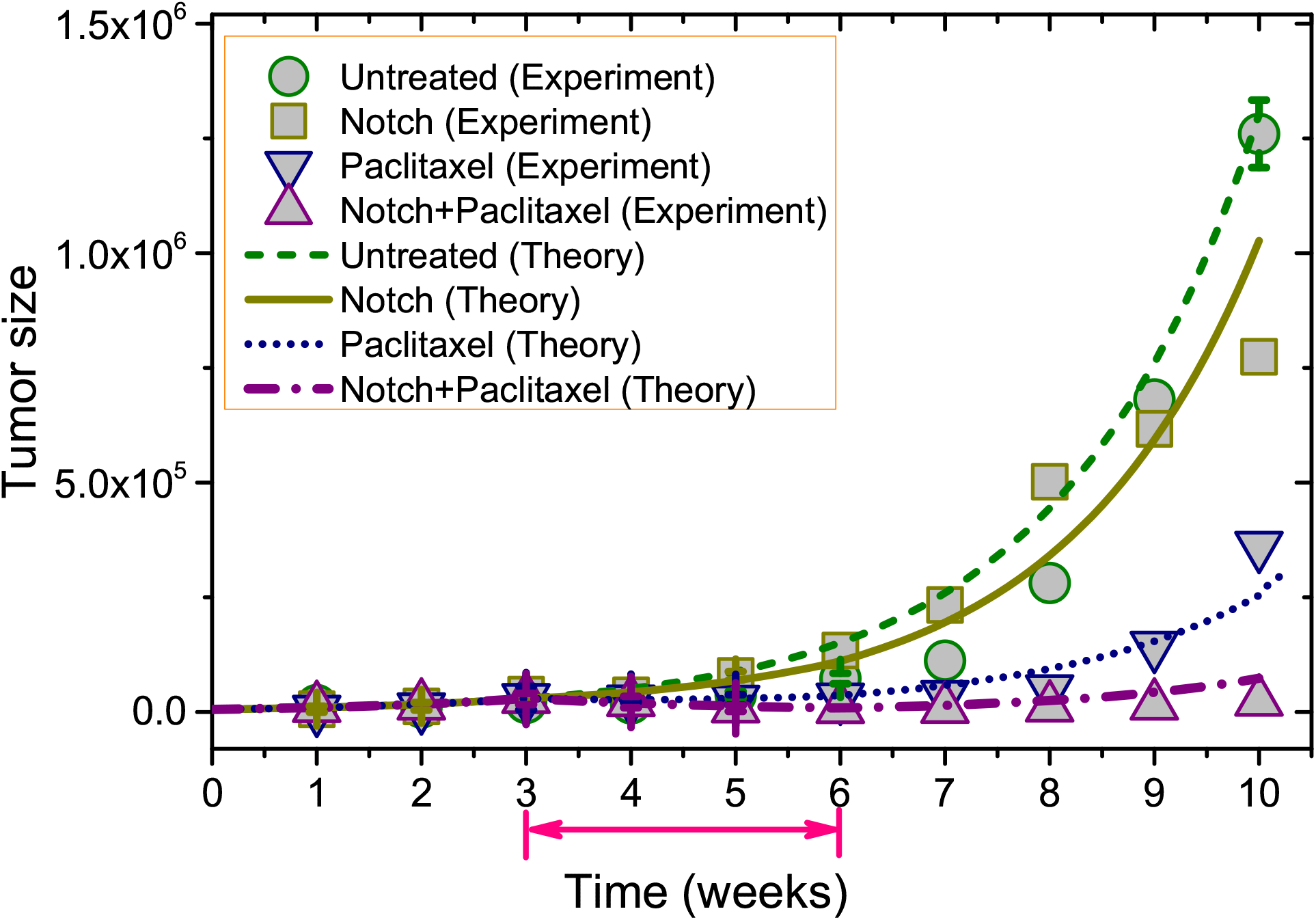
The dynamics of tumor growth under different conditions. The symbols represent results extracted from a recent experiment under four conditions^33^: The green circle shows the growth of mammary xenografts generated from parental circulating tumor cells (a mixture of HER2+ and HER2− cells) of breast cancer patients without any drugs. The dark yellow square and blue down triangle illustrate the growth of mammary xenografts under treatment of Notch inhibitor (*γ*-secretase inhibitor) and Paclitaxel from the 3rd to the 6th week (indicated by the double-headed arrow), respectively. The violet up triangle corresponds to the growth of mammary xenografts under treatment of both drugs simultaneously in the same period of time. The theoretical predictions for tumor growth under the four different cases are shown by the lines. The tumor is imaged using IVIS Lumina II. Its size is in the unit of the photon flux, which is proportional to the number of tumor cells.

Parameter values that are similar to the ones used to describe the experimental results *in vitro* are used but with the least changes for all the parameters in order to capture the tumor growth observed in *in vivo* experiments. By changing a rescaled factor *β* for *K*_1_ and *K*_2_ to estimate the rates (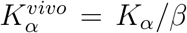, with *α* = 1 or 2) *in vivo, we can calculate the tumor (untreated) growth as a function of time and compare it to the experimental results. We find the optimum value β* ≈ 2.06*, which captures the tumor growth behavior in the in vivo experiments (see the green circles and dashed line in Fig. 3). The tumor seems to grow slower in vivo compared to in vitro (β* > 1*), which could be caused by the different imaging methods, or spatial constraint from other tissues, extracellular matrix, nutrient supply, etc*^42^.

HER2+ cells are sensitive to cytotoxic/oxidative stress (such as Paclitaxel treatment) while the HER2− cell shows a negligible response to Paclitaxel. On the other hand, Notch and DNA damage pathways are activated in the HER2− cell leading to sensitivity to Notch inhibition. However, the HER2+ cells are resistant to drugs for Notch inhibition^33^. To assess the influence of drugs on tumor growth, we set the effective growth rate 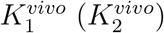 of symmetric cell division to a value, *γ*, when the drug, Paclitaxel (Notch inhibitor), is utilized during treatment. Then, we calculate the tumor size as a function of time under the treatment of both drugs and compared it to the experimental results. We find the best value *γ* ≈ −0.5, which captures the experimental observation for tumor growth (see the dash-dotted line in Fig. 3). We did not change the values of the asymmetric division rate constants, *K*_12_ and *K*_21_, due to the small value and also for simplicity.

Following the experimental protocol, we first let the tumor grow from a parental circulating tumor cells (78% of HER2+ and 22% of HER2− cells) with an initial size taken at week one. We then mimicked drug treatment from the third week to the sixth week. Surprisingly, our theory describes the growth dynamics of the heterogeneous tumor for different drug treatments well (see the different lines in Fig. 3). Our model successfully captured the inhibition of tumor growth under Paclitaxel treatment. Also the weak response of tumor under the treatment of Notch inhibitor emerges from our model naturally.

To understand the three distinct responses of the tumors to the drug treatments, shown in Fig. 3 further, we computed the time dependence of the tumor size in the first six weeks derived from our model with the treatment of either Notch inhibitor or Paclitaxel (see Figs. S2a and S2b in the SI). The tumor continues to grow rapidly without showing any clear response when treated with Notch inhibitor (see the symbols in navy in Fig. S2a), inhibiting the growth of HER2− cells. Although unexpected, the observed response can be explained from the cellular composition of the tumor. The fraction of HER2+ cells is high (> 70%) before drug treatment, and it increases monotonically to even higher values (~ 90%) during treatment, as shown in Fig. S2c in the SI. Considering the proliferation rate of HER2+ cells is higher than HER2− cells, it is clear that tumor response under Notch inhibitor only targets a minority of the tumor cell population and its reduction can be quickly replenished by the rapid growth of HER2+ (see the simple illustration in Fig. S2e under the treatment of Notch inhibitor).

Such a weak response is explained directly from the mean fitness, the growth rate *ω* = (*K*_1_ + *K*_12_)*f*_1_ + (*K*_2_ + *K*_21_)*f*_2_, landscape of the population, shown in Fig. 4. Without treatment, the mean fitness *ω* has a large constant value (see the dotted and solid lines in Figs. **4a** and 4**b**), indicating that the tumor grows at a steady rate aggressively. On the other hand, a relatively large initial value (*ω* ~ 0.3, see the location of yellow parts of the dotted or solid lines in Figs. **4c** and 4**d**) is still found, which shows a continuous growing phase of the tumor when subject to Notch inhibitor treatment. Therefore, no clear response would be observed if this treatment protocol is used. In addition, the tumor becomes even more aggressive with time (see the increasing value of *ω* in Figs. 4**c** and 4**d**) until it reaches to a maximum rate close to the untreated case.

**Figure 4:**
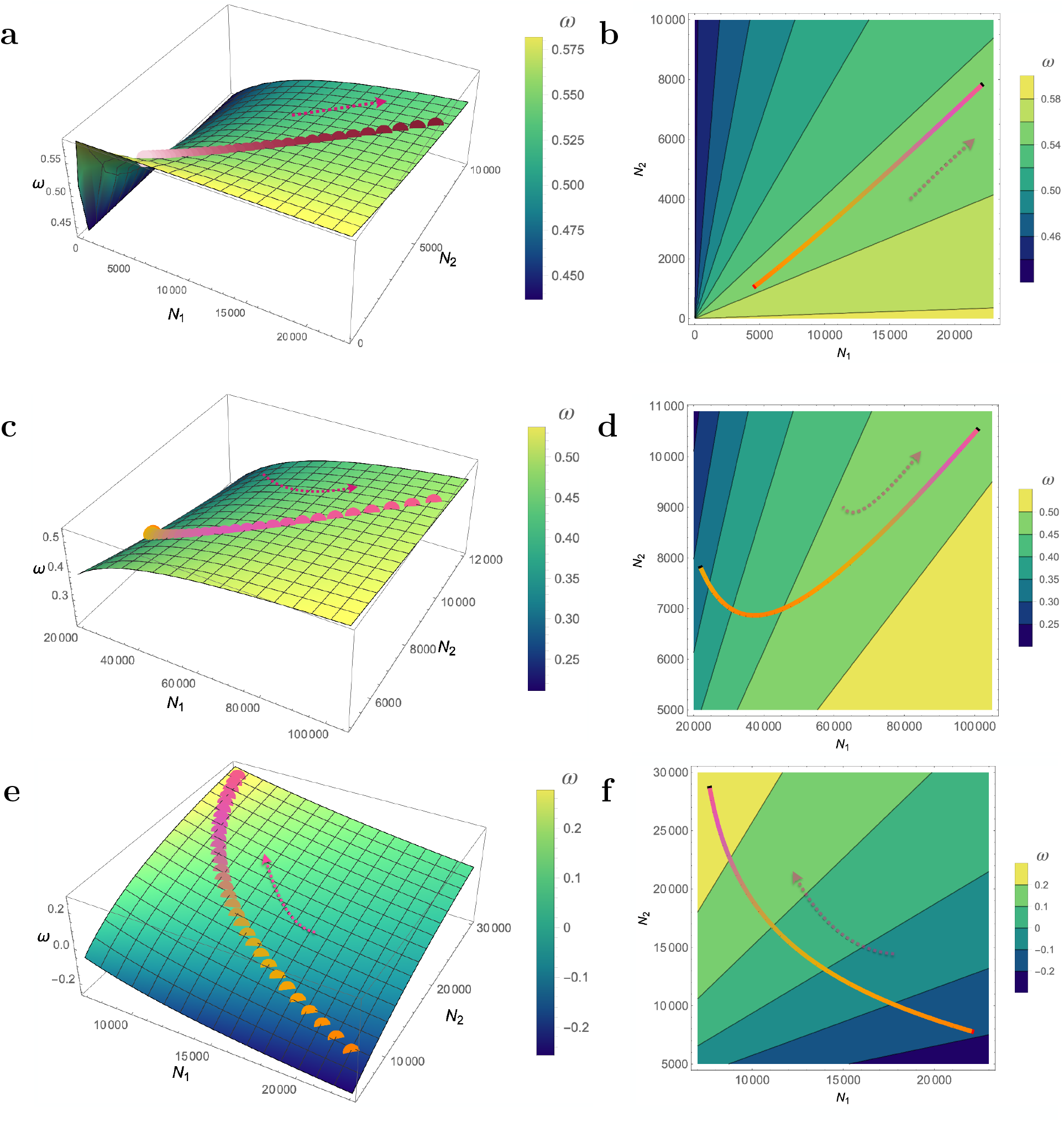
Fitness landscapes of cancer cell populations with and without drug administration. The mean fitness (*ω*) of the population as a function of the sizes (*N*_1_, *N*_2_) of the two subclones without any drugs (**a**) for the first three weeks, treated by Notch inhibitor (**c**) or Paclitaxel (**e**) from the 3rd to the 6th week, as shown in Fig. 3. The dotted lines show the phase trajectories for two cell populations along the fitness landscapes during treatment (all end with a larger total population size). The corresponding contour plots for **(a), (c), (e)** are shown in **(b), (d), (f)**, respectively with the phase trajectories indicated by the solid lines. The fitness *ω* (per week) is also colored by its value as indicated by the color bar on the right of each figure. The arrow indicates the direction of increasing time.

In contrast to the negligible effect of Notch inhibitor to the progression of the heterogeneous tumor, Paclitaxel treatment that targets the HER2+ cell leads to a clear reduction in the tumor size, and delays cancer recurrence (see Fig. S2b in the SI). Such a response is due to the high fraction of the HER2+ cell in the tumor. It leads to the slowly growing of HER2− cells, which cannot compensate for the quick loss of HER2+ cells at the start of the treatment (see the rapid decay of HER2+ cell fraction in Fig. S2d and Fig. S2e for illustration). However, the tumor recovers the fast growing phase in the fourth week (see Fig. S2b) after the drug is used, corresponding to the time when the fraction of HER2+ cell reaches around 0.5 (derived from our model with 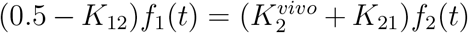, and see also Fig. S2d). Once the fraction of HER2+ cells decreases to small values, the proliferation of resistant HER2− cells can compensate for the loss of HER2+ cells. Just as discussed above, such a response can also be seen directly from the fitness landscape of the population under treatment of Paclitaxel (see Figs. 4**e** and 4**f**). The initial *ω* (~ −0.2) is negative during treatment (see the location of yellow parts of the dotted or solid lines in Figs. 4**e** and 4**f**), which indicates a shrinkage of the tumor. Such a state remained for some time until *ω* becomes positive. Although the value of *ω* increases with time, the tumor grows at a much lower rate at the end of Paclitaxel treatment compared to the situation when Notch inhibitor is used (see Figs. 4**c**–4**f**).

The fraction of HER2+ cells quickly recovers to the value in the stationary state after drug removal (see Figs. S2c and S2d), and the tumor grows aggressively again (see the insets in Fig. S2a-b and Fig. S2e for illustration). Therefore, the progression of the heterogeneous tumor cannot be controlled by a single drug, as demonstrated in the experiments, explained here quantitatively.

### Sequential treatment strategy

Our theory, and more importantly experiments, show that the utilization of two drugs simultaneously could significantly delay the recurrence of tumors compared to the treatments using only a single drug of either type (see Fig. 3). However, the quantity of drugs used in the former protocol is much higher than in the latter case. Also, both drugs (Paclitaxel and Notch inhibitor) have strong toxic side effects on normal tissues^43,44^. In the following, we consider a sequential treatment strategy with one drug followed by the treatment with the other, which would reduce the quantity of drugs used, and possibly reduce the toxic side effects.

In the sequential treatment, there are two alternative methods depending on the order in which the drugs are administered. We first let the tumor grows till the third week, and then apply the first drug, Notch inhibitor (Paclitaxel), from the third to the sixth week followed by the utilization of the second drug, Paclitaxel (Notch inhibitor), from the sixth to the ninth week. We used the same parameter values as taken in Fig. 3. Interestingly, we predict a dramatic difference between the responses of the tumors to the two treatment methods (see Fig 5**a**). The tumor size shows no clear response to the treatment by Notch inhibitor, increasing rapidly until Paclitaxel is used (see the circles in navy in Fig. 5**a** and a schematic illustration in the upper panel of Fig. 5**c**). From the phase trajectory (see the circles in Fig. 5**b**), a rapid increase of HER2+ cell population (*N*_1_) is found while HER2− cell population (*N*_2_) decays slowly. In contrast, just as shown in Fig. 3, a clear delay is observed for the tumor growth when treated with Paclitaxel first followed by Notch inhibitor (see the diamonds in pink and navy in Fig. **5a** and the lower panel of Fig. **5c** for illustration). Meanwhile, HER2+ and HER2− cell populations shrink rapidly during each drug treatment, as illustrated by the phase trajectory in Fig. 5**b** (see the diamonds). It indicates the effectiveness of these two drugs. In addition, the tumor size is always much smaller in the second protocol compared to the first, reaching three fold difference in size (see the tumor size at the sixth week in Fig. 5**a**). It follows that the order of drug administration greatly influences the treatment effects in the sequential treatment method, which is consistent with recent studies^45,46^. We also illustrate the tumor response when treated with the two drugs simultaneously (see the pentagons in Fig. 5**a**). A much better response is predicted compared to the treatment by Notch inhibitor first, followed by Paclitaxel (see the circles in Fig. 5**a**). However, the treatment by Paclitaxel first, followed by Notch inhibitor shows a similar good response with a close tumor burden at the end of treatment (see the diamond and pentagon in Fig. 5**a**). Hence, it is possible to find an optimal strategy to obtain a similar treatment effect with attenuated side effect.

**Figure 5:**
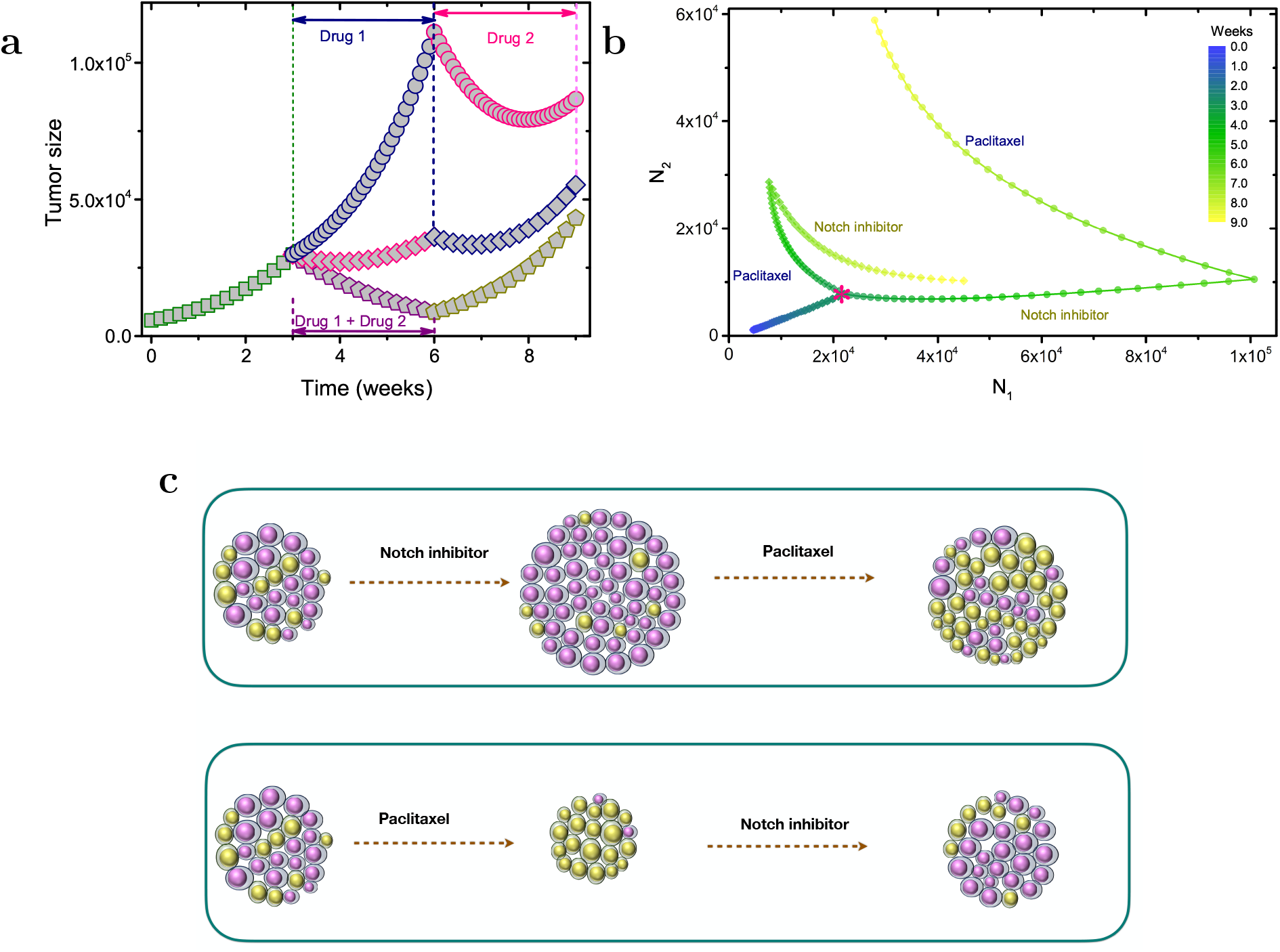
Tumor response using a sequential protocol for two drugs. (**a**) Comparison of drug responses for tumors under different treatments. The green squares show tumor growth before treatment. The tumor under the treatment of Notch inhibitor first (navy), then Paclitaxel (pink) is indicated by the circles. The diamonds show the tumor growth under the reverse order of drug treatment, Paclitaxel first (pink), followed by Notch inhibitor (navy). The pentagons demonstrate the treatment with both drugs administered simultaneously (violet color). The pentagons in yellow show the tumor growth after the removal of all drugs. The parameter values are the same as in Fig. 3. (**b**) The phase trajectories for the two subpopulations, HER2+ (*N*_1_), HER2− (*N*_2_), under two sequential treatments considered in Fig. 5**a**, respectively. The same symbols (circle and diamond) are used in (**a**) and (**b**). The initiation of the drug treatment is indicated by the red star and the trajectory color indicating the time is shown by the color bar. The drug name during each treatment period is also listed in the figure. (**c**) Illustration of the tumor responses under a sequential treatment of two drugs in different orders.

### Effect of duration of treatment

In the previous sections, a futile treatment with rapid tumor growth is frequently found (see Fig. 3 or the data in Figs. 5**a**–5**b**). We surmise that one drug should be removed at an appropriate time once it produces no benefits. We studied the influence of treatment period length (*τ*_*d*_) on tumor responses. First, we investigated the sequential treatment by Notch inhibitor followed by Paclitaxel for different *τ*_*d*_ values (see Fig. 6**a**). The phase trajectories show that the variations in *N*_1_, and *N*_2_ and their maximum values become smaller as *τ*_*d*_ is shortened. In addition, the response for each drug treatment is strengthened and the total tumor size (see the inset in Fig. 6**a**) is always smaller for a smaller *τ*_*d*_. Therefore, a small *τ*_*d*_ should be used when such a treatment method is applied.

**Figure 6:**
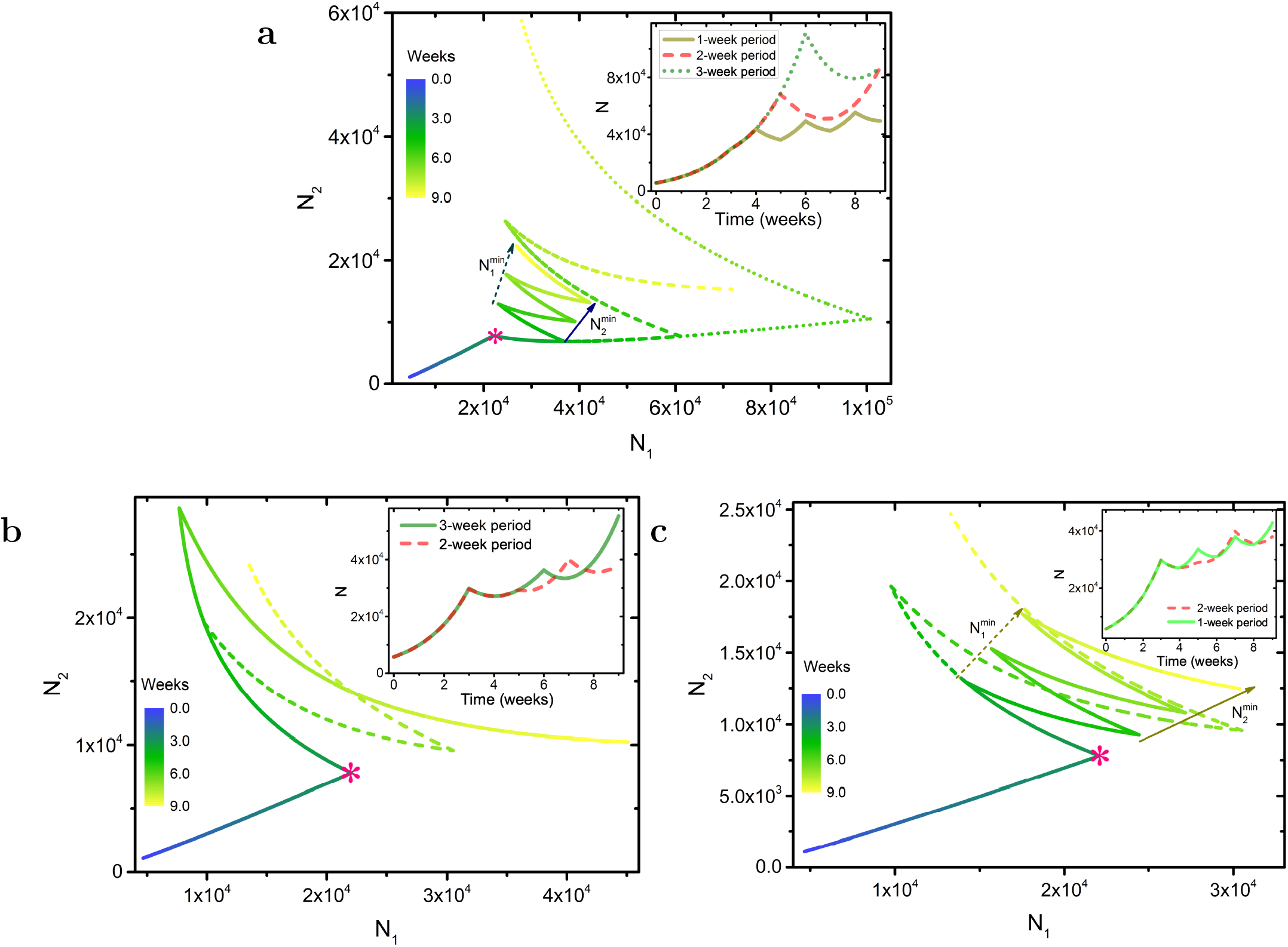
Phase trajectories for the two subpopulations under two different treatments. (**a**) Same as Fig. 5b, treated by Paclitaxel first, followed by Notch inhibitor. The treatment period (*τ*_*d*_) for each drug is one (solid line), two (dashed line) and three weeks (dotted line), respectively. (**b**) Same as Fig. 6**a** but the order of treatment is reversed (Paclitaxel first, then Notch inhibitor). The value of *τ*_*d*_ for each drug is three (solid line), and two weeks (dashed line), respectively. (**c**) Same as Fig. 6**b**, treated by Paclitaxel first, then Notch inhibitor. The treatment period for each drug is two (dashed line), and one week (solid line), respectively. The inset shows the total number (*N = N*_1_ + *N*_2_) of tumor cells as a function of time for each treatment.

Next, we performed a similar analysis for the treatment with Paclitaxel first, followed by Notch inhibitor (see Figs. 6**b**–6**c**). In contrast to the situation described above, the variations for *N*_1_, *N*_2_, and their responses to each drug treatment are similar even as *τ*_*d*_ varies. However, the total population size (see the inset in Figs. 6**b**–6**c**) is smaller for the two-week treatment compared to three and one-week treatment. We surmise that instead of using one-week treatment for each drug, a two-week period would be a better choice in this treatment strategy. Fig. 6 shows that the minimum values of 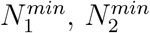 (see Figs. 6**a** and 6**c**) and the total minimum tumor size *N*^*min*^ (see the inset in Fig. 6) at each treatment cycle increases with time, irrespective of the value of *τ*_*d*_. This would result in uncontrolled tumor growth. In the following sections, we will discuss potential approaches to control the tumor burden even if it cannot be fully eradicated.

### Control of tumor burden

Despite the good response through certain treatment protocols as discussed above, tumor suppression is only transient, and the tumor recurs sooner or later due to drug resistance. Nevertheless, we can still seek, at least theoretically, a stable tumor burden as a compromise, which is similar to the goals of adaptive therapy^47^. For the breast circulating tumor cells consisting of HER2+ and HER2− cells, the model suggests that it is possible to control the tumor maintained at a constant size (with relatively small variations) (see Fig. 7**a**). Using a sequential treatment strategy, with Paclitaxel first, followed by Notch inhibitor and repeating the procedure periodically, the tumor burden may be kept at bay as it was before any treatment. The order of drug administration is important, as described above. During the treatment, it is efficacious to target the larger cell subpopulation with one specific drug until it becomes the minority, and then treat with the second drug. The quantity of drugs during each treatment should also be tuned to inhibit the growth of HER2− cells more efficiently (see the different values of 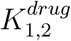 used in Fig. 7**a**). The time periods during the treatment of the two drugs are quite different. According to theory, a longer period of treatment is required for HER2+ cells (around two weeks as discussed in the previous section, see also Figs. 6**b**’6**c**) compared to the HER2− cells (around one week as discussed above, see also Fig. 6**a**). From the phase trajectory shown in Fig. 7**b**, a “limit cycle”-like structure is found in which the two subpopulations are well-controlled, and almost return to their original values after each round of treatment. In addition, we also found that the plasticity of breast cancer cells influences the tumor response during treatment. It appears that the tumor may be controlled or even eliminated eventually if we can inhibit the cellular plasticity by regulating related pathways such as EZH2, and Notch^34^ (see Fig. S3 and the detailed discussions in the Supplementary Information). Therefore, theoretical models based on the tumor evolutionary process are likely to be useful in predicting the tumor progression, the clinical response, and possibly in designing better strategies for cancer therapy^48–52^.

**Figure 7:**
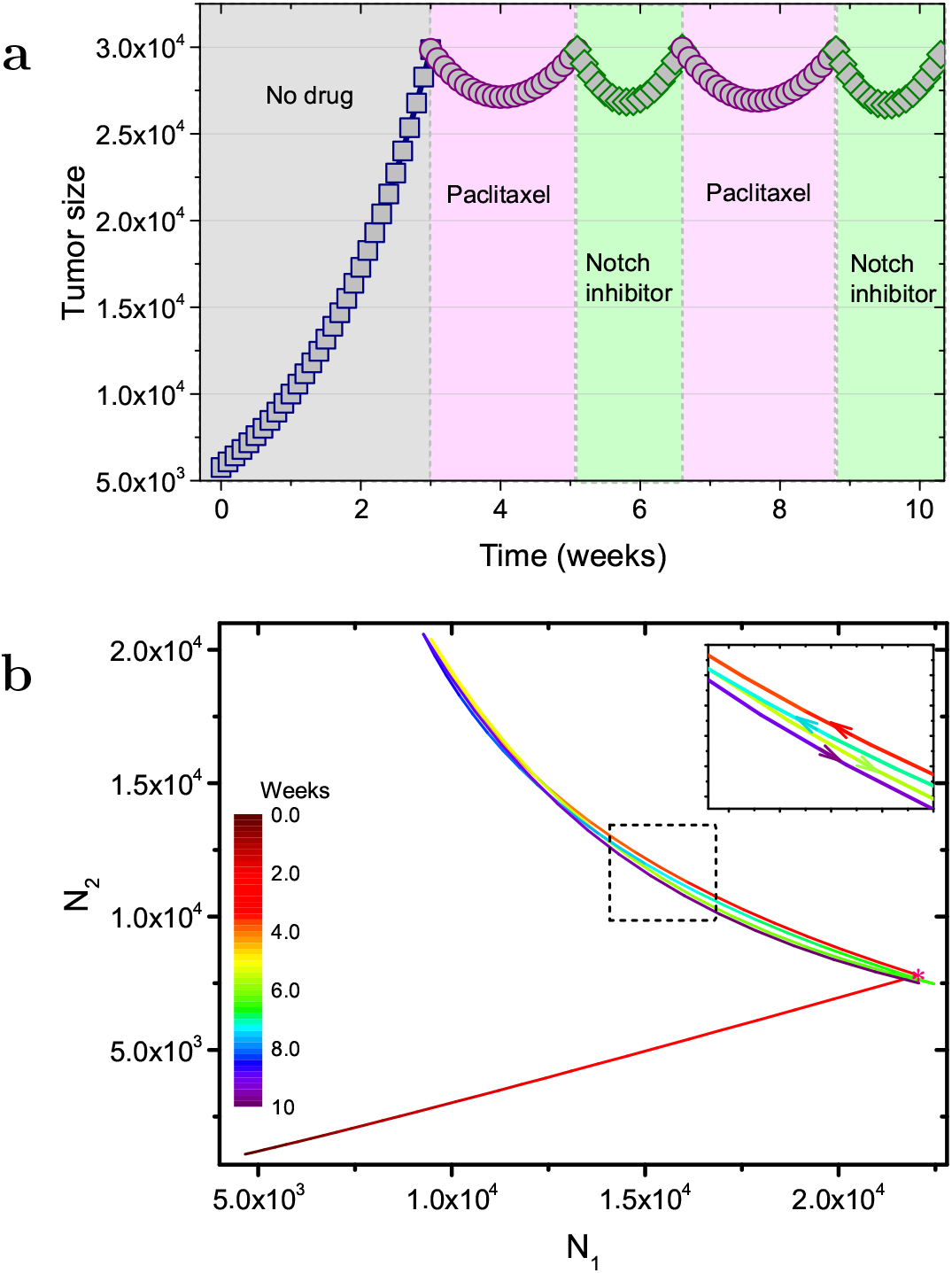
Tumor size control through a sequential treatment. (**a**) The Paclitaxel (violet regions) and Notch inhibitor (green regions) are used alternatively during the treatment. The parameter 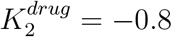. Other parameters are same as in Fig. 3. (**b**) Instead of the total population size, the phase trajectory for the two subpopulations is shown and a “limit cycle”-like structure is found under the treatment in Fig. 7**a**. The limit cycle is vividly illustrated in the inset, which shows a zoom-in of the dashed rectangle. The time increases from red, green, cyan to violet color as indicated by the arrows and the color bar on the left of the figure.

## DISCUSSION

We investigated the emergence of intratumor heterogeneity in breast cancer arising from cellular plasticity, which is embodied in the conversion between the HER2+ and HER2− phenotypes. In contrast to the unidirectional differentiation of normal stem cells^53,54^, many cancer cells demonstrate a great degree of plasticity that results in reversible transitions between different phenotypes, leading to intratumor heterogeneity without genetic mutations^28,31^. Such transitions are frequently observed in rapidly growing tumors, which is often neglected in theoretical models^28^. Although some studies have recognized the need for taking a growing population, the models typically have many unknown parameters^32,55^, which are hard to interpret.

By introducing a direct coupling between cell division and transition between phenotypes into a theoretical model, we provide a quantitative explanation for the emergence of a stable intratumor heterogeneity, a hallmark in HER-negative breast cancer patients. Our model accurately describes the evolution of different cancer cell fractions, and also the total tumor size observed in a recent single-cell experiment successfully. We predicted that the symmetric cell division appears more frequently compared to the asymmetric case for both types of cells found in breast circulating tumor cells. Using the same parameter values derived from the tumor growth experiments, our prediction for the cell fraction as a function of the cell colony size agrees well with experimental results. The cell colony size (5~8 cells) calculated from our theory for the emergence of one cell phenotype from the other is in good agreement with the experimental observations (5~9 cells).

The asymmetric cell division has not been observed in the breast circulating tumor cell experiment directly, although the experiment implies that cells of one phenotype produce daughters of the other phenotype^33^. However, in a more recent experiment this was detected in breast cancer^34^. It was found that the newly formed cell doublet, after one cell division, can be the same cell type (symmetric division) or different (asymmetric division, producing two daughter cells with one expressing the cytokeratin K14 while the other does not). It is also possible that the state transition is not only coupled to cell division but can also appear through tumor microenvironment remodeling^56^. However, inclusion of these processes will add two more free parameters to our model, which is not needed to give the agreement between theory and experiments. In addition, such a state transition is not observed after cytokinesis was inhibited in breast cancer experiment^34^. Nevertheless, our mathematical model could be extended to incorporate these possibilities should this be warranted in the future.

Although the asymmetric cell division explains the bidirectional state transition, the underlying mechanism for such an asymmetric division is still unclear. In the experiments^28,33,34^, the different states of cancer cells are mainly determined by the expression level of one or several proteins. It is possible that these proteins (HER2, K14, etc.) are redistributed in the daughter cells unequally during cell division, which could be realized through a stochastic process or regulation of other proteins^34,57,58^.

The reversible phenotype transitions in cells have been found in many different types of cancers^59–61^, which not only lead to the development of drug resistance but also induce very complex drug responses, as discussed here. Although each cell type is sensitive to one specific drug, the heterogeneous tumor derived from breast circulating tumor cells shows an obvious response to Paclitaxel but not to Notch inhibitor. Our model provides a quantitative explanation for the different time courses of the tumor under distinct treatments. The failure of the Notch inhibitor, even at the initial treatment is due to its target, HER2− cell which is a minority in the heterogeneous cell population, and has a slower proliferation rate compared to the HER2+ cell. Both experiments and our theory show a significant delay of tumor recurrence under the combination treatment with two drugs applied to the tumor simultaneously. We also predict that a sequential treatment strategy with Paclitaxel first, followed by Notch inhibitor (not in a reverse order of drugs) can show similar treatment effect as the one with two drugs used at the same time. In addition, the sequential treatment reduces the quantity of drugs administered each time, which can reduce the adverse effects and selection pressure for cancer cells from the drugs in principle^45^. Although not reported in the experiments^33^ and discussed in our present model, treatment-induced mutations in cancer cells have been reported for different chemotherapeutic drugs^62,63^. These new mutations may promote additional escape mechanisms and lead to treatment failure, which could be investigated in further studies.

One advantage of the mathematical model is that we can steer the evolutionary dynamics of each subpopulation by applying the right drug at the appropriate time to control the tumor burden. This allows for a fuller exploration of the parameter space, which cannot be easily done in experiments. Finally, we propose that patients could benefit from drugs which inhibit the plasticity of the cancer cells^34^. Taken together, our model could be applied to explore intratumor heterogeneity found in other type of cancers^34,59–61^. From the examples presented here and similar successful studies, we expect that the physical and mathematical models may provide a quantitative understanding for the cancer progression and also stimulate new ideas in oncology research^19,48,64–66^. We should emphasize that mathematical models sharpen the questions surrounding the mechanisms of intratumor heterogeneity, but real data from patients are needed to understand the origins of intratumor heterogeneity.

## Acknowledgements

We are grateful to Shaon Chakrabarti, Abdul N Malmi-Kakkada, and Sumit Sinha for discussions and comments on the manuscript. This work is supported by the National Science Foundation (PHY 17-08128), and the Collie-Welch Chair through the Welch Foundation (F-0019).

## Author contributions

X.L. and D.T. conceived and designed the project, and co-wrote the paper. X.L. performed the research.

