## Supplementary Information for "A mathematical model for phenotypic heterogeneity in breast cancer with implications for therapeutic strategies"

Xin Li[1, a)] and D. Thirumalai[1, b)]

[1]*Department of Chemistry, University of Texas, Austin, TX 78712, USA.*

(Dated: 16 October 2021)

[a)]Electronic mail:

[b)]Electronic mail:

In the Supplementary Information (SI), we provide several additional figures, and related discussions that are pertinent to the results in the main text.

**The tumor growth dynamics from a single cell type:** Fig. S1 shows the proliferation dynamics of FACS-purified HER2+ (green circles) and HER2- (blue triangles) cell subpopulations from cultured circulating tumor cells, respectively. It can be noted that the proliferation rate for HER2+ is faster than for HER2- cell, as also observed from the proliferation marker (Ki67)[1]. In contrast, there is no difference in the apoptotic rates for the two cell types from the apoptotic markers (cleaved-caspase 3 or annexin 5)[1]. This is consistent with the results in Fig. S1, and our theoretical calculations with $K_1 - K_2 = \Sigma \approx 0.3 > 0$. The solid lines in Fig. S1 are derived from Eqs. (1)-(2) in the main text from the expression $N(t) = N_1(t) + N_2(t)$ with one free parameter $K_2 \approx 0.7$, and by implication $K_1 \approx 1.0$.

**Drug response for a heterogeneous cancer cell population:** Fig. S2 shows our theoretical calculations for the drug response of a tumor cell population generated from a mixture of HER2+ (78%) and HER2- (22%) cells under treatment of Notch inhibitor (see Fig. S2**a**) or Paclitaxel (see Fig. S2**b**) from the 3rd to the 6th week. There are no drugs for the first three weeks, which results in a rapid growth of the cell population. Figs. S2**c** and S2**d** show the dynamics of the cell fraction ($f_1(t)$) for HER2+ cells during and after the treatment of drugs. The Fig. S2**e** is an illustration of the tumor responses during and after the treatment with the two drugs.

**Cellular plasticity leads to failure of treatments:** We have learned from our calculations that the plasticity of breast cancer cells is one of the leading reasons for ITH, which in turn leads to drug resistance during therapy. We investigated ITH influences the tumor response during treatment further. By varying the values of $K_0$ ($\equiv K_{12} = K_{21}$), which could vary among patients, we found that a strong transition between the two cell states can lead to total failure of treatments (see Fig. S3**a**). The two cell subpopulations and the total tumor size show a much weaker response during each drug treatment, irrespective of the order of drug administration (see Fig. 5a in the main text and Fig. S3**a**). This suggests that enhanced cellular plasticity could lead to an easy escape of cancer cells from the targeted drugs. On the other hand, we found that it is much easier to control the tumor burden as the cellular plasticity is inhibited (see Fig. S3**b**), which leads to a strong tumor response when treated with the two drugs. Surprisingly, the total tumor size is similar at the end of the two treatments with different order of administration of the two drugs, although protocol that uses paclitaxel first is still a better choice. It appears that it may be achieved by controlling the tumor burden (see Fig. S3**c**) or even eliminating the tumor eventually if we can inhibit the cellular plasticity by regulating related pathways such as EZH2, and Notch[2]. The model shows that if $K_{12} = K_{21} = 0$, then ITH, which is a cause of drug resistance, disappears starting from either type of tumor cells. In this theoretical scenario, cancer may be eradicated by administering the appropriate drugs, even if it is a heterogeneous cell population before treatment.

### SUPPLEMENTARY FIGURES

Figure SI. **The time course of the population size initiated from a single cell type**

Figure S2. **The dynamics of tumor growth under different drugs**

Figure S3. **Targeting cellular plasticity**

### SUPPLEMENTARY TABLES

Table 1. **Model parameters**

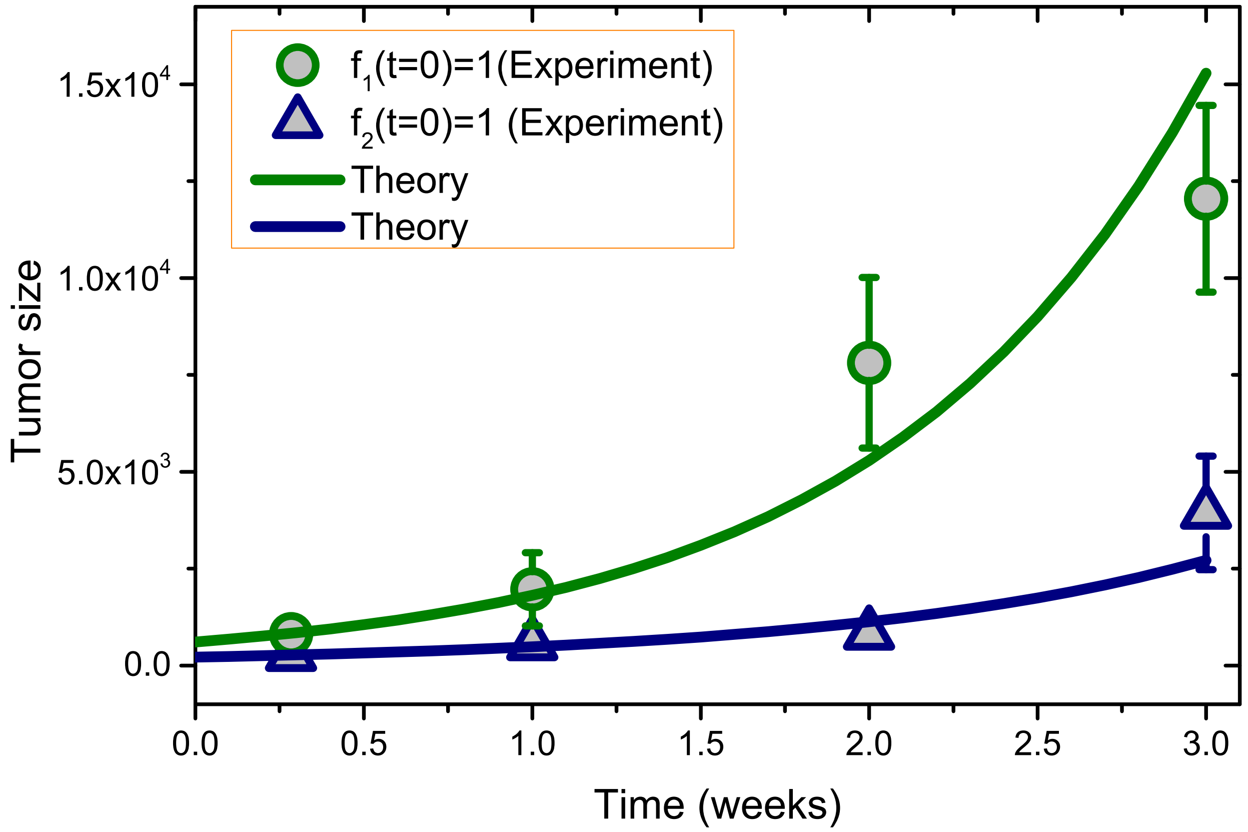


**Figure** S1: **The time course of the population size initiated from a single cell type.** The cell population size as a function of time initiated from only HER2+ (green) or HER2- (navy) circulating tumor cells. The symbols represent experimental results[1] and the solid lines correspond to theoretical results. The cell population is imaged using CellTiter-Glo assay. Its size is in the unit of luminescence count which is proportional to the tumor cell number.

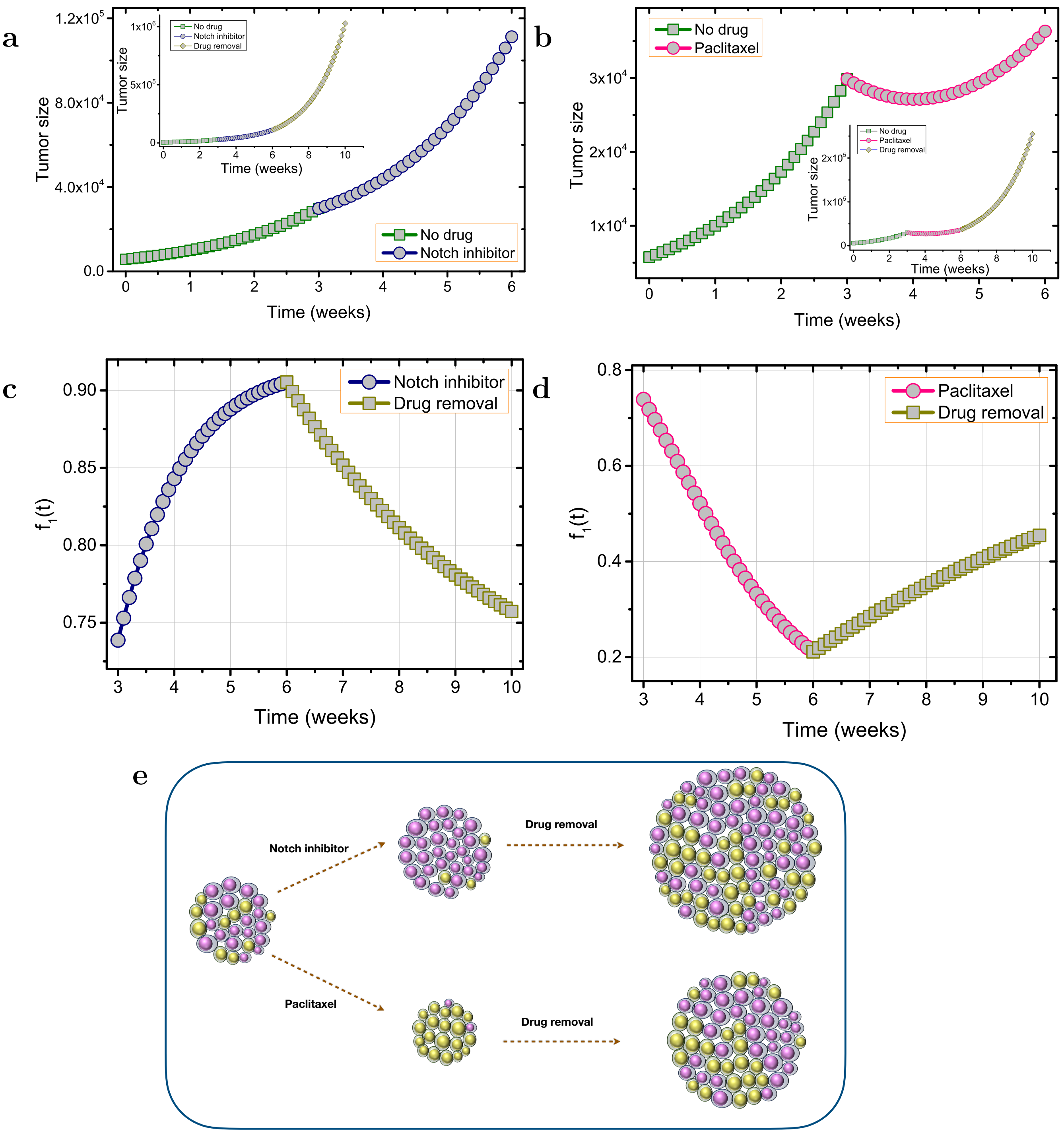


**Figure** S2: **The dynamics of tumor growth treated with different drugs.** Same as Fig. 1 in the main text, the tumor size as a function of time under Notch inhibitor (**a**) or Paclitaxel (**b**) treatment from the 3rd to the 6th week. The insets in (**a**) and (**b**) also include the time evolution of the tumor size after drug removal. The fraction ($f_1(t)$) of HER2+ cells as a function of time subject to treatment of Notch inhibitor (**c**) and Paclitaxel (**d**), respectively. (**e**) Illustration of the tumor responses under two different treatments and the development of drug resistance.

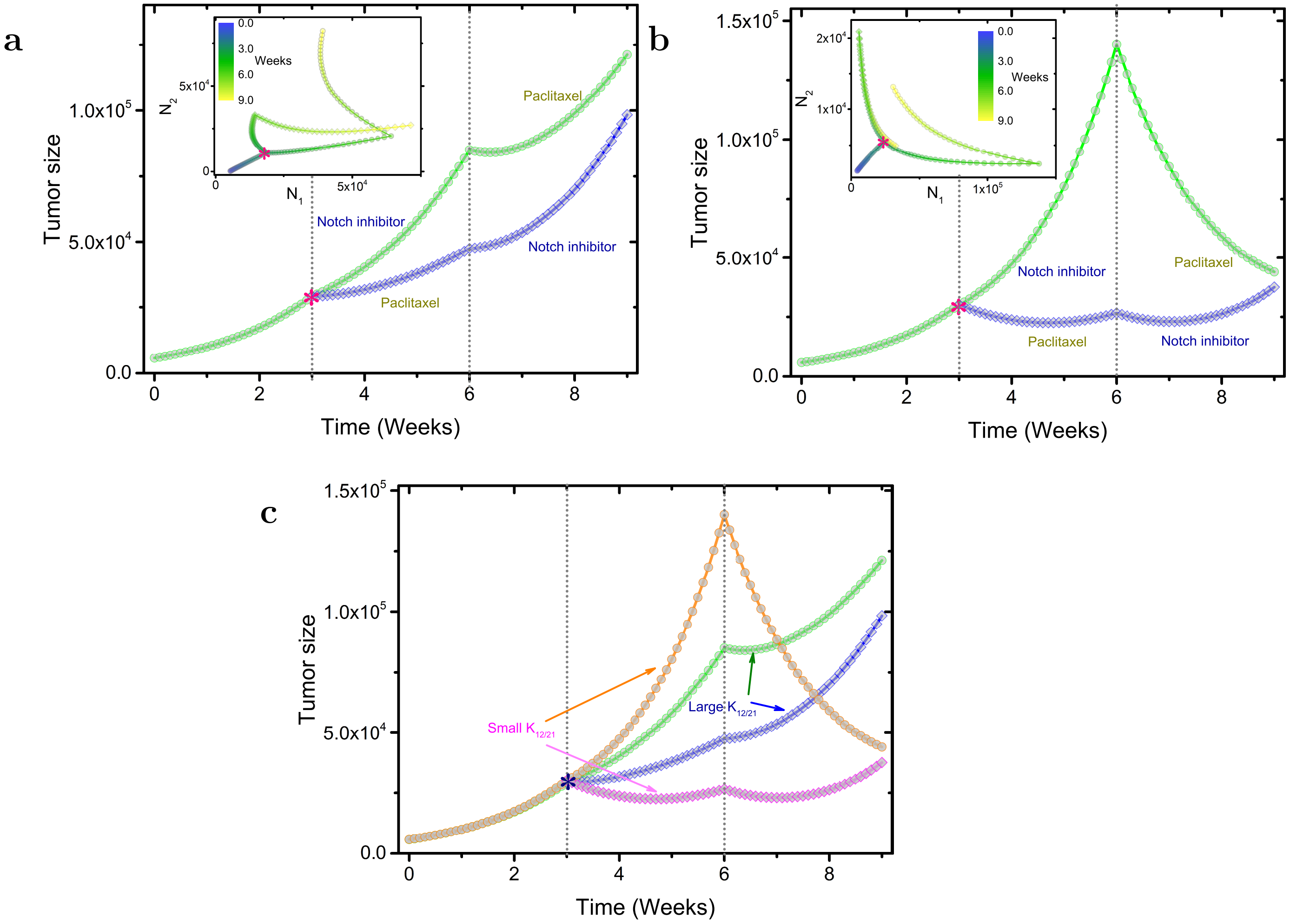


**Figure** S3: **Targeting cellular plasticity** (**a**) Tumor response for a sequential treatment. Same as Fig. 5a in the main text, except for a larger asymmetric division rate ($K_{12} = K_{21} = 5K_0 = 0.45$) for the two cell types. The inset shows the phase trajectory for the two subpopulations obtained by different order in which the drugs are administered. Treatment by giving Paclitaxel first, then Notch inhibitor is shown as diamonds, and the circles show treatment by Notch inhibitor first followed by Paclitaxel. (**b**) Same as Fig. **S3a** except for a smaller asymmetric division rate ($K_{12} = K_{21} = 0.1K_0 = 0.009$) for the two cell types. The total cell division rate for each cell ($K_1 + K_{12}, K_2 + K_{21}$) and also other parameters are the same as in Fig. 3 in the main text. (**c**) Comparison of the tumor responses subject to two treatment methods with different values of asymmetric division rates shown in Figs. **S3a** - **S3b**.

TABLE I: The parameters used in the model. The parameter values with "*" are only used in Fig. S3 to investigate how the treatment effect could be influenced by the cellular plasticity, which can be quite different among patients. Other values are derived from comparison between theory and experiments[1].

| **Parameters** | **Values** |
|---|---|
| $K_1$ | 1.0 |
| $K_2$ | 0.7 |
| $K_{12} = K_{21}$ | 0.09, 0.009*, 0.45* |
| $K_1^{vivo}$ | 0.49 |
| $K_2^{vivo}$ | 0.34 |
